# Synaptic active zones are ordered nanostructures designed by supramolecular block copolymers

**DOI:** 10.64898/2025.12.15.691964

**Authors:** Hirokazu Sakamoto, Shunsuke F. Shimobayashi

## Abstract

Neurotransmitters are released at presynaptic active zones, where a conserved cytomatrix exhibits nanoscale order first visualized by electron microscopy over half a century ago. However, the physical principles underlying this architecture have remained unclear. Here we show that active zone scaffolds act as protein block copolymers, whose nanoscale microphase separation produces hexagonally packed nanodot arrays. Super-resolution imaging combined with block-copolymer theory identifies ELKS and RIM as the minimal pair sufficient to reproduce this ordered lattice, mediated by orthogonal interactions between ELKS coiled-coil domains and RIM intrinsically disordered regions. Additional active zone proteins, including RIMBP and Liprin-α, modulate lattice spacing and morphology in a stoichiometry-dependent manner, coupling nanoscale order to neurotransmission. More broadly, block-copolymer microphase separation emerges as a design principle for constructing functional nanostructures in living systems.

---

Synaptic transmission underlies information processing in the brain, relying on the temporally precise release of neurotransmitters at presynaptic terminals. At these sites, specialized active zones ensure that synaptic vesicles fuse in close proximity to voltage-gated calcium channels, enabling rapid and reliable signal transfer with nanoscale spatial precision and millisecond temporal accuracy (*1, 2*). Since they were first visualized in electron micrographs in the 1960s, active zones have been recognized as containing electron-dense protein assemblies with a periodic organization of approximately 100 nm (*3, 4*). This supramolecular “cytomatrix” is conserved across species and comprises multiple scaffolding proteins, including ELKS, RIM, RIMBP, Liprin-α, and Munc13, which collectively arrange vesicles and channels into ordered nanoscale architectures (*1, 2, 5*). Despite this long-standing knowledge of their ultrastructure and molecular components, the biophysical logic by which these proteins self-organize into such nanoscale order has remained elusive. In this study, we show that active zone scaffolds, exemplified by ELKS and RIM, behave as supramolecular block copolymers that self-organize into ordered nanolattices, thereby uncovering the physical principle that underlies nanoscale organization of presynaptic active zones.

## Results

### ELKS2–RIM1 as the minimal pair of active zone proteins generating ordered nanolattices

To dissect the molecular basis of active zone nanostructure formation, we focused on two core scaffold proteins, ELKS2 (also known as CAST) and RIM1 (Figure 1A) (*6–8*). Genetic deletion of both ELKS and RIM family genes severely disrupts active zone architecture and abolishes synaptic vesicle docking and release (*9*). ELKS2 is composed primarily of three coiled-coil domains (CC1–CC3), whereas RIM1 contains extensive intrinsically disordered regions (see Methods, Figures 1B and 1C). We first examined their native organization by immunolabeling rat brain sections (Figure 1D). Using confocal laser scanning microscopy (CLSM) at diffraction-limited resolution, ELKS2 and RIM1 signals overlap almost completely at cortical synapses (Figure 1E). Stimulated emission depletion (STED) microscopy then revealed that ELKS2 organizes into punctate nanoclusters of 62.3 ± 14.2 nm (Figures 1F and 1G), consistent with previous EM observations of active zone materials (*1–4, 10, 11*).

**Figure 1.**
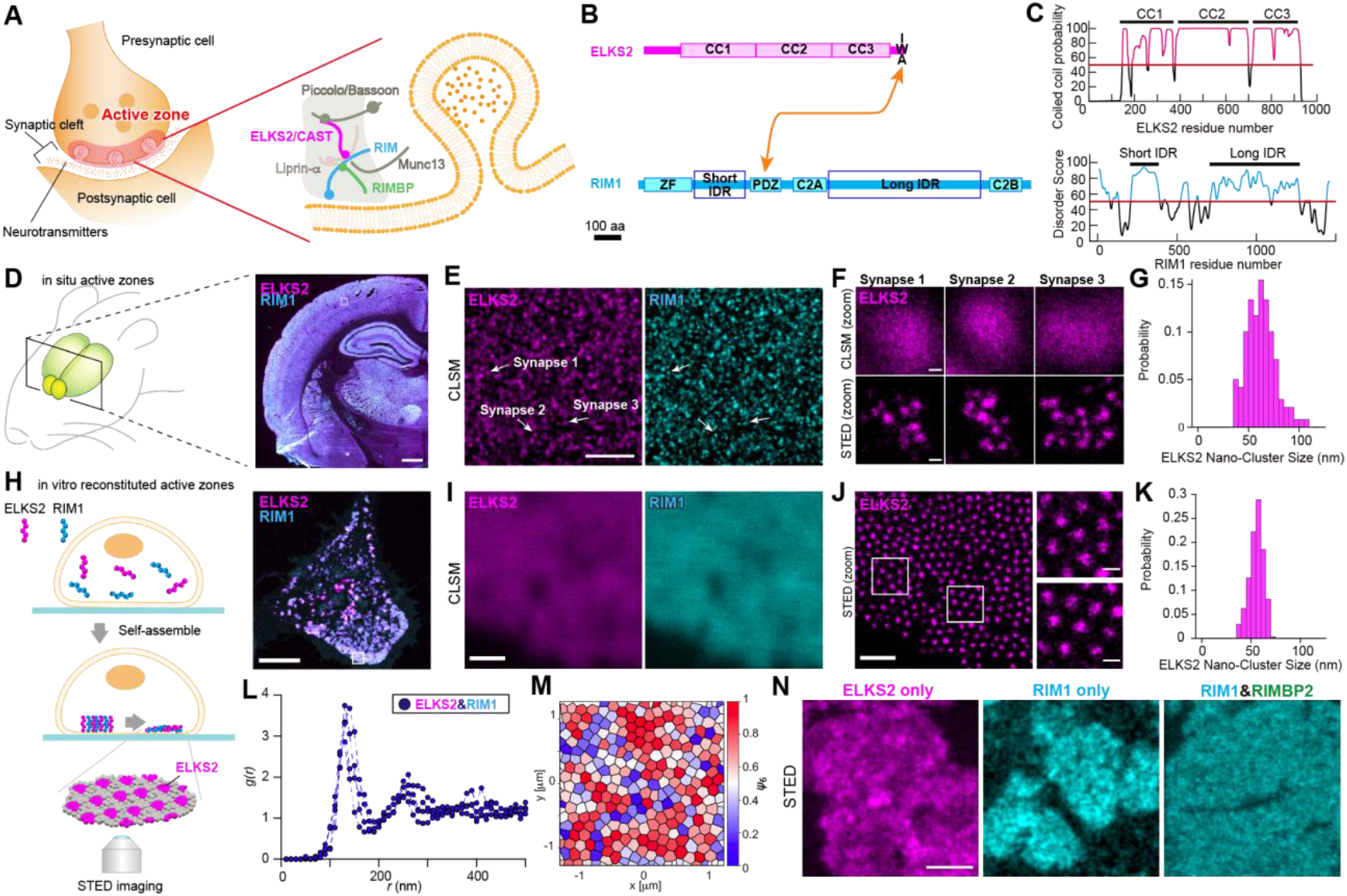
ELKS2–RIM1 as the minimal active zone module forming ordered nanolattices. (**A**) Schematic of presynaptic active zones. (**B**) Domain organization of active-zone components ELKS2 and RIM1. (**C**) Coiled-coil (CC) and intrinsically disordered region (IDR) predictions for ELKS2 and RIM1. (**D**) In situ presynaptic active zones in a rat brain tissue section, stained for ELKS2 and RIM1. Scale bar: 1 mm. (**E**) Enlarged confocal laser scanning microscopy (CLSM) images of the white-boxed region in (D). Synapses are identified as punctate sites of overlapping ELKS2 and RIM1 immunofluorescence. Scale bar: 5 mm. (**F**) STED (stimulated emission depletion) images of the same synapses as in (E), revealing nano-clustered ELKS2. Scale bar: 100 nm. (**G**) Probability distribution of ELKS2 nanocluster size for in situ active zones with *n*_dots_ = 239 in 20 synapses. (**H**) Schematic of in vitro reconstitution (left) and a representative CLSM image of non-neuronal COS-7 cells (right). Co-expression of SNAP-ELKS2 (magenta) and RIM1-EGFP (cyan) on the plasma membrane drives self-assembly into ELKS2–RIM1 condensates. Scale bar: 8.5 mm. (**I**) Enlarged CLSM images of the white-boxed region in (H). Scale bar: 5 mm. (**J**) STED imaging of the region shown in (I): overview (left) and two zoomed views of the boxed regions (right). Scale bar: 1 mm (left), 200 nm (right). (**K**) Probability distribution of ELKS2 nano-cluster size in vitro with *n*_dots_=270. (**L**) Radial pair-correlation function *g(r)* of nanostructures in reconstituted AZs, showing a primary peak at ∼ 130 nm, indicating characteristic nano-cluster spacing *n*_cells_ =4, *n*_dots_ =3394. (**M**) Voronoi diagrams derived from a representative STED image of ELKS2–RIM1 nanodot condensates on the COS-7 cell membrane, with the hexatic order parameter *ψ*_6_ visualized using color coding. (**N**) STED images of condensates on the plasma membrane of COS-7 cells expressing ELKS2 only, RIM1 only, or RIM1 together with RIMBP2. Scale bar: 500 nm. *n*_cells_: number of cells. *n*_dots_: total number of nanodots.

To test whether ELKS2 and RIM1 alone can recapitulate active zone nanostructure, we co-expressed SNAP-ELKS2, fused to a membrane translocation signal, together with RIM1-EGFP on the plasma membrane of non-neuronal COS-7 cells, where endogenous expression of ELKS and RIM is negligible (Figure 1H). Although no periodic clustering was apparent by CLSM, STED imaging of these condensates revealed regularly spaced ELKS2 nanoclusters of 55.4 ± 7.3 nm diameter (Figures 1I–1K), closely matching the size in rat synapses. The two-dimensional radial distribution functions, *g(r)*, exhibited multiple peaks— indicative of a repeating lattice—with the first peak at 132.0 ± 4.5 nm (see Methods, Figure 1L), consistent with earlier electron microscopy measurements (*10*). To characterize angular symmetry within these lattices, we computed the bond-orientational order parameter, which revealed a clear sixfold symmetry (*ψ*_6_)— analogous to that seen in hexagonally packed colloidal monolayers, block-copolymer thin films, and even in biological tissues (Figures 1M and S1; see Methods) (*12, 13*). Expression of membrane-anchored ELKS2 or RIM1 alone produced membrane-associated condensates without periodic order, and co-expression of RIM1 with RIMBP2 likewise failed to generate nanodot arrays (Figure 1N). Thus, ELKS2 together with RIM1 define the minimal module sufficient to assemble the ordered nanostructure characteristic of presynaptic active zones.

### ELKS2–RIM1 act as supramolecular protein block copolymers that self-assemble into hexagonal nanolattices

To characterize how ELKS2 and RIM1 organize at the nanoscale, we imaged both ELKS2 and RIM1 by STED microscopy in COS-7 cells, revealing ELKS2 as punctate “dot” clusters and RIM1 as complementary “ring” structures with ∼130 nm spacing (Figure 2A). Notably, this spatial segregation reflects the distinct molecular localizations of ELKS2 and RIM1 (Figure 2B). Using condensate recruitment assays in human osteosarcoma (U2OS) cells (see Methods, Figure 2C), where overexpression induces ELKS2 or RIM1 condensates, we quantified homotypic interactions by measuring partition coefficients *K* and corresponding Gibbs free-energy changes Δ*G* (see Figure 2D and Methods): ELKS2 self-associates strongly with Δ*G* ≈ −3.1 *k*_B_*T*, and RIM1 similarly with *ΔG* ≈ −2.1 *k*_B_*T*, each capable of independently forming condensates (Figures 2E and F). By contrast, their high-affinity heterotypic binding is mediated by the C-terminal IWA motif of ELKS2 and the PDZ domain of RIM1, with a dissociation constant *K*_d_ ≈ 0.3 µM (*14, 15*) (Figure 2G). To further probe heterotypic contacts between ELKS2 coiled coil domains and RIM1 intrinsically disordered regions, we analyzed recruitment within ELKS2 condensates: RIM1, which does not form condensates on its own at moderate expression levels, is fully recruited with Δ*G* ≈ −3.0 *k*_B_*T*, whereas deletion of either IWA or PDZ abolishes this recruitment (Figures 2D, 2H and 2I). These data reveal that ELKS2 and RIM1 assemble as a supramolecular diblock-copolymer—two homotypic blocks bridged by high-affinity IWA–PDZ binding—driving the robust formation of sixfold symmetric nanodot arrays in living cells.

**Figure 2.**
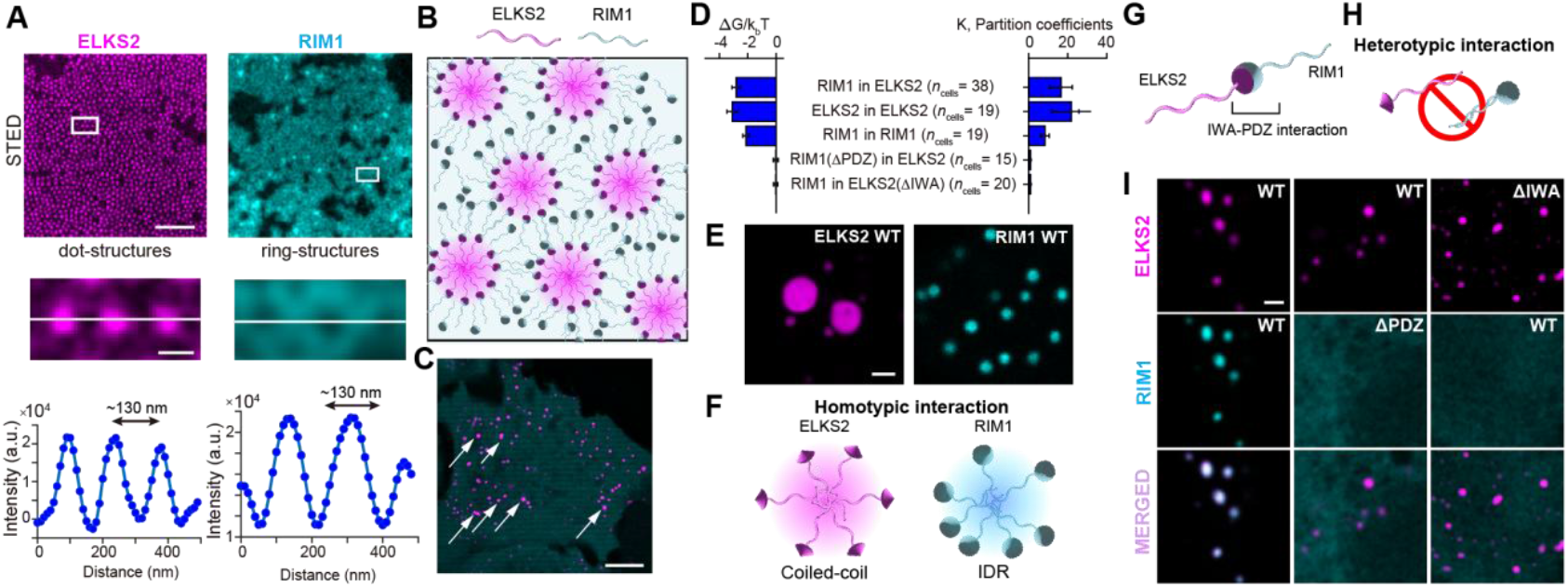
ELKS2–RIM1 act as supramolecular protein block copolymers self-assembling into hexagonal lattices. (**A**) STED images of COS-7 cells expressing ELKS2 (magenta) and RIM1 (cyan) tagged with SNAP (top); enlarged views of the white-boxed regions showing ELKS2 dot-structures and RIM1 ring-structures (middle); fluorescence intensity profiles along the white lines in the middle panels (bottom), revealing periodic peaks separated by ∼ 130 nm. Scale bars: 1 mm (top), 100 nm (middle). (**B**) Schematic of the supramolecular block-copolymer model of ELKS2 and RIM1 driving periodic assembly. (**C**) CLSM images of human U2OS cells co-expressing mGFP-ELKS2 (magenta) and mCherry (cyan), showing punctate ELKS2 condensates at the plasma membrane indicated by white arrows. Scale bar: 10 mm. (**D**) Bar graphs of the dimensionless chemical potential difference Δ*G/k*_B_T (left) and partition coefficient *K* (right) measured for ELKS2 and RIM1 condensates under WT and mutant conditions. (**E**) CLSM images of U2OS cells expressing mGFP-ELKS2(WT) (left) or RIM1(WT)-mRuby3 (right), demonstrating homotypic condensate formation. Scale bar: 2 mm. (**F**) Schematic of homotypic interaction mechanisms for ELKS2 (via coiled-coil domains) and RIM1 (via intrinsically disordered regions). (**G**) Schematic of supramolecular block-copolymer assembly via IWA–PDZ interaction between ELKS2 and RIM1. (**H**) Schematic of negligible heterotypic interaction between ELKS2 and RIM1 in the absence of IWA–PDZ binding. (**I**) CLSM images of U2OS cells co-expressing mGFP-ELKS2 and RIM1-mRuby3 under three conditions: WT ELKS2 with WT RIM1 (left), WT ELKS2 with RIM1(ΔPDZ) (middle), and ELKS2(ΔIWA) with WT RIM1 (right). Note that RIM1 alone does not form condensates at moderate expression levels. Scale bar: 2 mm. *n*_cells_: number of cells.

### Block-copolymer physics governs the organization of ELKS2–RIM1 nanostructures

Block copolymers undergo microphase separation into rich, periodic nanostructures—lamellae, gyroids, and hexagonally packed cylinders—a cornerstone of soft-matter and materials science (*16*). To test whether ELKS2–RIM1 assemblies obey similar block-copolymer physics, we first constructed a covalently linked ELKS2–RIM1(ΔN) “block copolymer” in silico and in vitro (Figures 3A and B), bridged at the IWA-PDZ interaction site. In silico, we modeled ELKS2–RIM1 as a covalent diblock copolymer—two flexible blocks of length *N*/2—using the Ohta–Kawasaki functional (*17*), which predicts hexagonal, lamellar, and coexistence morphologies as a function of the interaction parameter *χN* and ELKS2 area fraction *f*_ELKS2_ (Figure 3A; see Methods). Experimentally, a single-chain, covalent ELKS2–RIM1(ΔN) fusion formed condensates on the plasma membrane in COS-7 cells. STED imaging showed that these condensates assembled into hexagonal nanocluster arrays with ∼130 nm center-to-center spacing, matching those generated by supramolecular ELKS2–RIM1 assemblies (Figures 1J, 3C, and 3D).

**Figure 3.**
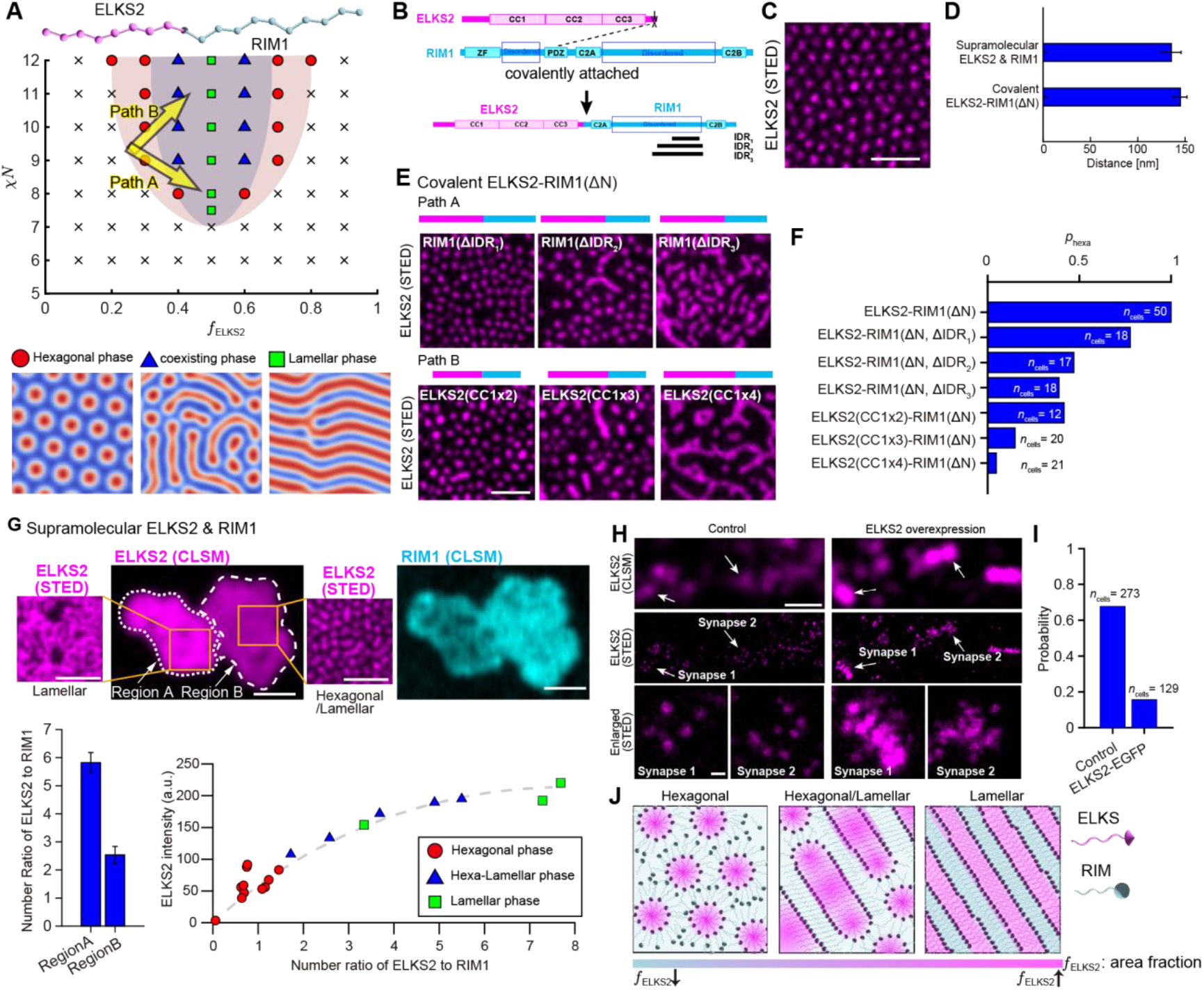
Block-copolymer physics governing ELKS2–RIM1 nanostructure organization. (**A**) Numerically determined phase diagram of ELKS2–RIM1(ΔN) diblock copolymer in two-dimensional simulations, plotted as functions of ELKS2 area fraction *f*_ELKS2_ and segregation parameter c*N*. Three phases—hexagonal, lamellar, and coexisting—are observed. (**B**) Schematic of covalently attached ELKS2 and RIM1(ΔN) constructs used to lock the supramolecular architecture. (**C**) STED image of a COS-7 cell expressing SNAP-ELKS2-RIM1(ΔN)-EGFP. Scale bar: 500 nm. (**D**) Periodic distance of nanostructures measured in COS-7 cells expressing either both ELKS2 and RIM1 (*n*_cells_ =11) or covalent ELKS2–RIM1(ΔN) (*n*_cells_ =11), showing similar spacing. (**E**) STED images of COS-7 cells expressing SNAP-ELKS2– RIM1(ΔN)–EGFP constructs with progressive RIM1 IDR deletions (Path A: ΔIDR_1_, ΔIDR_2_, ΔIDR_3_) or ELKS2 CC1 tandem repeats (Path B: CC1×2, CC1×3, CC1×4), corresponding to simulation paths A and B in (A), showing the nanoscopic transition from hexagonal to lamellar phase. Scale bar: 500 nm. (**F**) Probability of adopting the hexagonal phase, *p*_hexa_, for various ELKS2–RIM1(ΔN) mutants (*n*_cells_ =12–50). (**G**) Top, CLSM and STED images of COS-7 cells co-expressing SNAP-ELKS2 (magenta) and RIM1-EGFP (cyan), with dashed outlines marking two adjacent condensates that differ in ELKS2:RIM1 ratio and exhibit distinct nanostructures. Scale bar: 1 mm for CLSM images, 500 nm for STED images. Bottom left, bar graph of the ELKS2:RIM1 number ratio in Region A versus Region B. Bottom right, scatter plot of ELKS2 mean intensity per condensate versus ELKS2:RIM1 number ratio, demonstrating an apparent transition from hexagonal to lamellar-like phases, with the dashed line as a visual guide (*n*_cells_ =19). (**H**) CLSM (top row) and STED (middle row) images of primary hippocampal neuron active zones labeled with anti-ELKS2 antibody in control (left) and EGFP-ELKS2 overexpression (right) conditions, with enlarged STED views of the indicated synapses (bottom row; white arrows). Scale bars: 1 mm (CLSM), 100 nm (STED). (**I**) Probability of observing dot-structured nanoclusters exclusively in single synapses under control (*n*_synapses_ =273) and EGFP-ELKS2 overexpression conditions (*n*_synapses_ =129). (**J**) Schematic illustrating nanostructure transitions of ELKS2–RIM1 assemblies as a function of ELKS2 area fraction *f*_ELKS2_. *n*_cells_: number of cells. *n*_synapses_: number of synapses.

Based on in silico phase diagrams predicting a hexagonal-to-lamellar transition (Figure 3A), we hypothesized that raising *f*_ELKS2_ would drive the same switch in the experiments. To test this, we systematically elevated *f*_ELKS2_ via two complementary strategies—Path A (progressive deletion of RIM1 IDRs: ΔIDR1, ΔIDR2, ΔIDR3); and Path B (tandem duplication of the ELKS2 CC1 domain: CC1×2, CC1×3, CC1×4) (Figures 3A and B). Assuming that ELKS2 and RIM1 together fill the two-dimensional condensate area, deletion of RIM1 IDRs reduces the effective area contribution of RIM1 and thereby increases the relative area fraction of ELKS2, whereas CC1 tandem duplication directly enlarges the ELKS2 contribution. In each case, STED imaging revealed a continuous progression from pure hexagonal arrays, through coexistence of hexagonal and lamellar regions, to predominantly lamellar-like nanostructures, accompanied by a monotonic decrease in *p*_hexa_ —the fraction of cells exhibiting purely hexagonal dot order (Figures 3E and F). These results quantitatively validate our block copolymer theory predictions and confirm that *f*_ELKS2_ is the key parameter governing active zone nanostructure morphology.

In actual active zones, supramolecular ELKS2–RIM1 condensates rely on dynamic, noncovalent interactions whose local stoichiometry can fluctuate. To test whether the ELKS2:RIM1 ratio governs internal nanostructure, we imaged two adjacent condensates (Regions A and B) by STED microscopy (Figure 3G) and measured their ELKS2:RIM1 ratios using our covalent ELKS2–RIM1(ΔN) standard (Figure S2). Region A, with higher *f*_ELKS2_, apparently adopted a lamellar pattern, whereas Region B, at lower *f*_ELKS2_, exhibited hexagonal/lamellar coexistence (Figure 3G), in precise agreement with block-copolymer theory (*16*). Quantitative analysis confirmed that incremental increases in the ELKS2:RIM1 ratio shift condensates from hexagonal to coexistence to fully lamellar phases. Finally, to probe physiological relevance, we overexpressed EGFP–ELKS2 in primary hippocampal neurons and found that elevating ELKS2 levels within active zones converted synaptic nanostructures from punctate hexagonal arrays to connected, lamellar-like arrangements (Figures 3H and 3I), demonstrating that block-copolymer microphase separation actively sculpts presynaptic nanostructure in living neurons. Together, these results provide quantitative, biophysical evidence that ELKS2–RIM1 form a supramolecular diblock copolymer whose stoichiometry-driven microphase separation into hexagonal and lamellar domains governs the structural modulation of presynaptic active zone nanostructure (Figure 3J).

### Modular design and client control of ELKS2–RIM1 nanostructures and synaptic function

Given the emerging picture that ELKS2–RIM1 condensates generate periodic nanostructures via supramolecular diblock-copolymer behavior, we next examined how molecular features of ELKS2 and RIM1 encode distinct roles in nanostructure formation. To delineate the molecular architecture of ELKS2, we combined AlphaFold structure prediction with Marcoil coiled-coil mapping, which consistently partitioned ELKS2 into three contiguous coiled-coil segments, CC1–CC3 (Figure 4A) (*18–20*). This approach clarified the boundaries of each coiled-coil segment and distinguished them from intervening intrinsically disordered regions, thereby providing a structural rationale for treating CC1–CC3 as discrete modules. By contrast, RIM1 contributes extensive intrinsically disordered regions (Figures 1C and S3). To assign each segment’s contribution, we generated ELKS2–RIM1(ΔN) fusions lacking CC1, CC2, or CC3 and imaged their condensates by STED microscopy (Figure 4B). CC1 deletion abolished all periodic clustering, marking CC1 as the nucleating scaffold. CC2 removal preserved dot formation but sharply reduced inter-dot spacing, whereas CC3 deletion left spacing intact (Figure 4C). Although CC3 does not influence spacing, single-particle tracking showed that CC3 deletion shifts dot dynamics from sub-diffusive (α ≈ 0.2) to near-free diffusion (α ≈ 1), yielding an MSD increase of more than tenfold (Figures 4D and S4). Together, these findings demonstrate that CC1 drives nanocluster nucleation, CC2 specifies periodic spacing, CC3 maintains dynamic stability, and RIM1 IDRs provide the flexible matrix—highlighting how the combined modular domains of ELKS2 and RIM1 underpin multiple, distinct roles in nanostructure formation.

**Figure 4.**
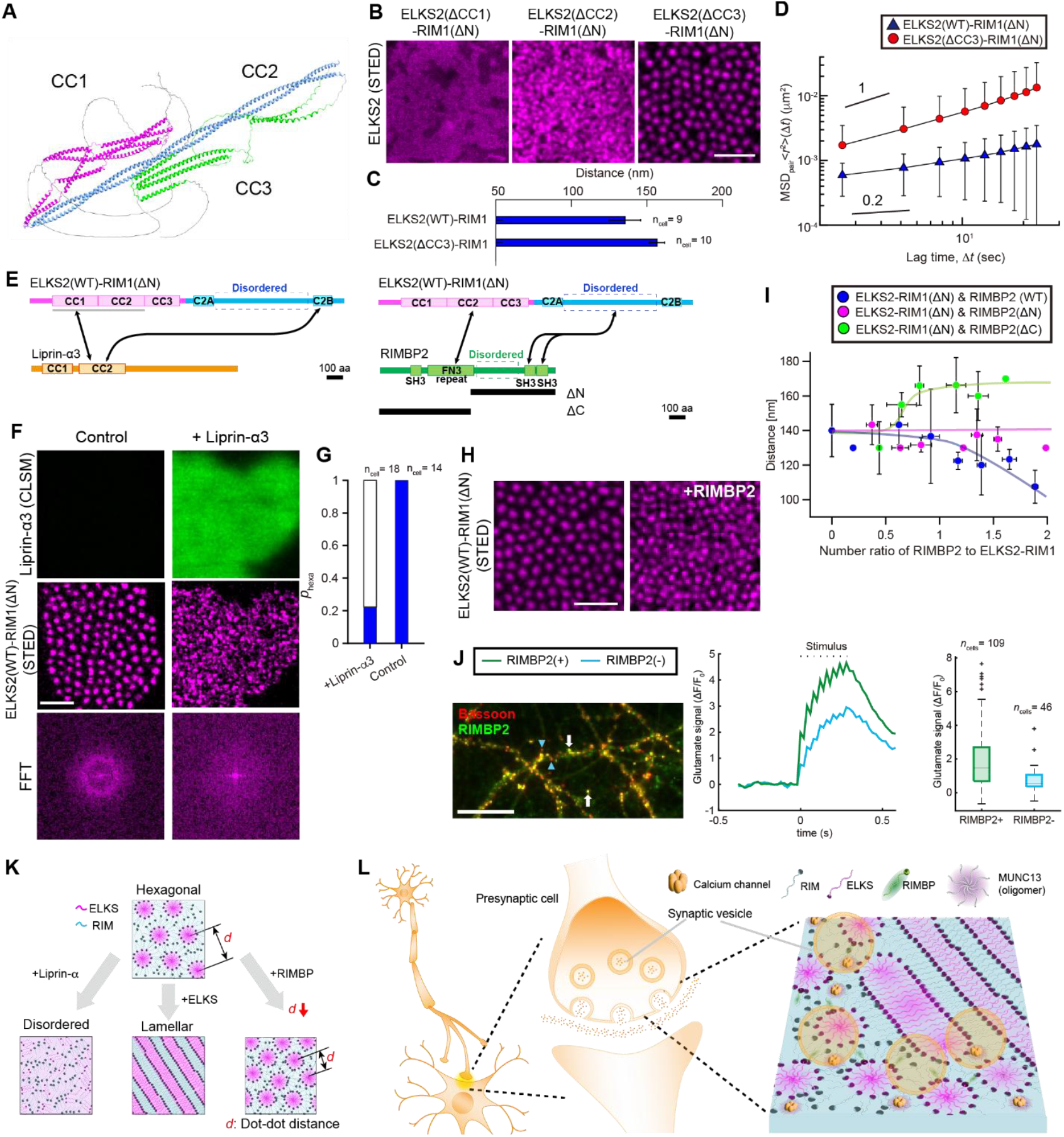
Modular architecture and client control of ELKS2–RIM1 nanostructures and synaptic output. (**A**) AlphaFold2-predicted ribbon model of the ELKS2 coiled-coil assembly, highlighting CC1 (magenta), CC2 (blue), and CC3 (green) domains. (**B**) STED images of COS-7 cells expressing ΔCC mutants of membrane-anchored SNAP-ELKS2–RIM1(ΔN)-EGFP, demonstrating CC-dependent nanostructure formation within condensates. Scale bar: 500 nm. (**C**) Comparison of periodic spacing between WT ELKS2– RIM1 and ΔCC3 assemblies, showing preservation of nanocluster periodicity. (**D**) Mean-square displacement (<Δ *r*^2^>) as a function of lag time Δ*t* for WT (*n*_cells_ = 8) and ΔCC3 ELKS2–RIM1 condensates (*n*_cells_ = 10), showing subdiffusive behavior for WT and normal diffusion for ΔCC3. Error bars indicate standard deviation. (**E**) Schematic of domain architectures for ELKS2–RIM1(ΔN) and Liprin-α3 (left); the arrows indicate interactions between Liprin-α3 CC2 and ELKS2 CC1/CC2 or RIM1 C2B domain. Schematics of ELKS2(WT)–RIM1(ΔN) and RIMBP2 constructs (right): RIMBP2 shown as full-length, N-terminal truncation (ΔN), and C-terminal truncation (ΔC); the arrows indicate interactions between ELKS2 CC2 and RIMBP2 FN3 repeats, and between RIM1 IDR and RIMBP2 SH3 domains. Scale bar: 100 aa. (**F**) CLSM images of Liprin-α3 (top row) and STED (middle row) images of ELKS2–RIM1 condensates in COS-7 cells under control and Liprin-α3-EGFP overexpression conditions, with FFT power spectra (bottom row) showing loss of hexagonal order upon Liprin-α3 overexpression. Scale bars: 500 nm. (**G**) Probability of hexagonal phase adoption, *p*_hexa_, in ELKS2–RIM1 condensates in the absence or presence of Liprin-α3. (**H**) STED images of ELKS2(WT)–RIM1(ΔN) condensates formed with or without co-expressed RIMBP2. Scale bar: 500 nm. (**I**) Plot of inter-cluster distance versus RIMBP2:ELKS2(WT)–RIM1(ΔN) ratio, showing construct-specific, dose-dependent modulation. (**J**) Immunofluorescence images of Bassoon and RIMBP2 in rat hippocampal neurons; white arrows and blue triangles denote RIMBP2-positive and RIMBP2-negative AZs, respectively. Right two panels: time trace data for stimulus-evoked glutamate release obtained by using a fluorescent glutamate sensor, eEOS (left; stimulation at 25 Hz, 8 APs) and corresponding quantification of peak signal amplitude (right). Green trace represents RIMBP2-positive active zones (*n*_synapses_=109); cyan trace represents RIMBP2-negative active zones (*n*_synapses_ =46). Scale bar: 2 mm. (**K**) Summary schematic of nanostructure phase behaviors: Liprin-α incorporation disrupts order, converting the lattice into a disordered state; increasing ELKS levels drive a hexagonal-to-lamellar transition; and RIMBP binding reduces dot-dot spacing *d*. (**L**) Model of presynaptic active-zone nanostructure regulation by ELKS– RIM assemblies and client proteins (RIMBP, Munc13, Calcium channels), coordinating vesicle docking and release.

Next, we sought to dissect how additional active zone clients tune this supramolecular architecture. Block-copolymer theory predicts that branching or bridging architectures alter the effective chain extension and thereby modulate inter-domain spacing *d*, either compacting or expanding the lattice depending on the symmetry of engagement (*21*). Prompted by this physical prediction, we examined how client factors such as Liprin-α3 and RIMBP2, which can bind both ELKS2 and RIM1 (*22, 35*) (Figures 4E and S5), regulate inter-dot spacing and supramolecular order. To address this, we co-expressed the covalent ELKS2– RIM1(ΔN) fusion with Liprin-α3 or RIMBP2 in COS-7 cells and confirmed client incorporation into condensates (Figures 4F and S5). Addition of Liprin-α3 to ELKS2–RIM1(ΔN) condensates converted the hexagonal array into a disordered state, as quantified by loss of sixfold FFT peaks and a dramatic decrease in *p*_hexa_ (Figure 4F and 4G). We speculate that the Liprin-α3 CC2 domain simultaneously engages the ELKS2 CC1/CC2 region and RIM1 C2B domain located at the end of its IDR, bringing these two scaffolds together at a single site and thereby abolishing the ordered nanostructure. In contrast, incorporating RIMBP2 into ELKS2–RIM1(ΔN) condensates progressively reduced dot-to-dot spacing as a function of the RIMBP2:ELKS2–RIM1 ratio, with a two-fold excess of RIMBP2 yielding a maximal contraction of ∼30 nm (Figures 4H and 4I). Given that RIMBP2 can bind both ELKS2 and RIM1 via its distinct domains, FN3 repeats and SH3 domains (Figures 4E, right; S5), we hypothesized that it bridges the two scaffold proteins to draw clusters closer together. To test this, we generated RIMBP2 truncations lacking either the N-terminal FN3 repeats (ΔN) or the C-terminal SH3 domains (ΔC), each predicted to engage only one partner. Neither truncation could compact spacing—in fact, the ΔC mutant showed a slight increase in spacing (Figure 4I), implying an effectively elongated chain extension of RIM1. This confirms that simultaneous dual engagement by full-length RIMBP2 is required for its spacing-compaction function. From the perspective of branched block copolymer theory, this dual engagement can be interpreted as creating a bridging circulation that reduces conformational freedom and thereby compacts the inter-dot spacing.

Moreover, to test the functional impact of RIMBP2-mediated spacing compaction, we examined the effect of RIMBP2 expression, visualized by immunocytochemical labeling, on glutamate release in rat hippocampal neurons (Figure 4J, left). Using the eEOS glutamate sensor (*5*), we recorded stimulus-evoked release at RIMBP2-positive (+) versus RIMBP2-negative (−) active zones and found that RIMBP2-positive sites exhibited significantly larger peak amplitudes (Figure 4J, right), indicating that tighter nanocluster spacing enhances glutamate release efficacy at presynaptic active zones. Together with the order-disrupting effect of Liprin-α3, these findings underscore that client factors dynamically sculpt the supramolecular nanostructure to fine-tune neurotransmitter release.

## Discussion

By integrating block-copolymer theory with super-resolution imaging of supramolecular ELKS2–RIM1 assemblies, we demonstrate that ELKS2–RIM1 nanoclusters obey classic microphase-separation physics to establish a hexagonal baseline architecture at presynaptic active zones, and that client factors dynamically optimize and control subsequent structural transitions and synaptic functions. While LLPS research has largely focused on IDR-mediated condensation, we show that ELKS2 coiled-coil domains (CC1–CC3) are essential for nanoscale ordering inside condensates, with distinct domains driving nucleation, spacing control, and dynamic stabilization. Moreover, we reveal diverse structural roles of active zone clients: elevated ELKS2 levels induce a hexagonal-to-lamellar transition that reshapes docking geometry; Liprin-α3, which is required for the assembly of active zones (*22*), disrupts the ordered nanostructure; and RIMBP2 compacts dot-to-dot spacing to boost release probability (Figure 4K). As one phase of the nanoscale microphase separations at active zones, RIM forms nanodomains that partner with Munc13 to organize vesicle release sites (*5*) and RIMBP to cluster VGCCs (*23*). This indicates that the nanoscale organization driven by ELKS and RIM is fundamental for controlling neurotransmitter release (Figure 4L).

Recently, a variety of intracellular, membraneless organelles—such as nucleoli and stress granules—have been shown to form via liquid–liquid phase separation (LLPS) (*24–28*). In synaptic compartments, LLPS has been implicated in multiple contexts (*29*): multivalent interactions between SynGAP and PSD-95 driving postsynaptic density (PSD) condensation in postsynaptic sites (*30*), synapsin-mediated clustering of synaptic vesicles into a dynamic “vesicle liquid phase” in presynaptic terminals (*31, 32*), and condensates of active zone scaffold proteins (Liprin-α/SYD-2, ELKS, RIM/RIMBP) that recruit calcium channels in presynaptic terminals (*33–36*). However, these studies have focused primarily on mesoscale condensates, without addressing the nanoscale architectures. Here, using super-resolution STED microscopy of ELKS2– RIM1 condensates on cellular membranes in living cells and mapping to block-copolymer phase diagrams, we reveal a previously overlooked paradigm—that nanoscale order within LLPS condensates, first glimpsed by electron microscopy in the 1960s (*3, 4*), is a fundamental driver of presynaptic active zone organization and functions. The discovered ELKS2–RIM1 nanodot lattice explains the molecular basis of the “dense projections” observed over six decades ago, setting the stage for revealing how nanoscale molecular order within LLPS condensates orchestrates precise vesicle and channel positioning for millisecond-precision neurotransmitter release. Our work bridges supramolecular chemistry, neuroscience, and materials science (*37, 38*), demonstrating how block copolymer physics, including its branched architectures, offers a unified framework for understanding and engineering functional protein nanostructures. This framework opens avenues for targeted synaptic modulation, programmable protein nanomaterials, and next-generation bio-inspired devices.

## Supporting information

Supplemental informaton

## Acknowledgments

We would like to thank Kenzo Hirose for fruitful discussions, Naoya Kimpara and Tetsuroh Ariyoshi for preliminary experiments, Masumi Sanada and Akira Miyaki for their experimental support, and Tomo Kurimura for assistance with figure preparation. We are also grateful to Yusuke Maeda (Kyoto University) and Tomofumi Yoshida (Kyoto University) for their critical reading of the manuscript. This work was supported by Japan Science and Technology Agency (JST) PRESTO (Grant Numbers JPMJPR21E7 to H.S. and JPMJPR21E8 to S.F.S.), the JST FOREST Program (Grant Numbers JPMJFR240X to H.S. and JPMJFR230V to S.F.S.), JSPS KAKENHI (Grant Numbers 19K16251, 21K15183, 25K02374, and 25H02354 to H.S.; 18K13521, 23K13072, and 25H01328 to S.F.S.), Japan Agency for Medical Research and Development (AMED) (Grant Number JP23bm1323001 to S.F.S.), iPS Cell Research Fund (to S.F.S.), Uehara Memorial Foundation (to S.F.S.), Nakatani Foundation (to S.F.S.), Kao Foundation for Arts and Sciences (to S.F.S.), iPS Academia Japan Grant 2022 (to S.F.S.), and Takeda Science Foundation (to S.F.S.).

## Author Contributions

Conceptualization, H.S. and S.F.S.; methodology, H.S. and S.F.S.; Investigation, H.S. and S.F.S.; writing— original draft, H.S. and S.F.S.; writing—review & editing, H.S. and S.F.S.; funding acquisition, H.S. and S.F.S.; resources, H.S. and S.F.S.; supervision, H.S. and S.F.S.

