## Supplemental informaton for "Synaptic active zones are ordered nanostructures designed by supramolecular block copolymers"

**This PDF file includes:**

Materials and Methods

Supplementary Text

Figures S1 to S5

Tables S1

[1]These authors contributed equally to this work.

### Materials and Methods

#### Animals

All animal experiments were performed in accordance with the policies of the Animal Ethics Committee of The University of Tokyo. Sprague-Dawley rats of either sex were commercially acquired from Japan SLC and housed in a group (2–4 per cage) with a 12 h light/dark cycle and food and water available ad libitum. Wooden or plastic toys and wood chip bedding were routinely provided.

#### Cell culture

COS–7, Glial, and 293T (for virus production only) cells were cultured in DMEM (FUJIFILM Wako Pure Chemical) supplemented with 10% FBS, 4 mM L-glutamine (FUJIFILM Wako Pure Chemical), 1 mM sodium pyruvate, and 1% penicillin/streptomycin at 37°C under 5% $CO_2$. U2OS and Lenti–X 293T (for virus production only) cells were maintained in Dulbecco's Modified Eagle's Medium (DMEM; Nacalai Tesque, #08458–16) supplemented with 10% fetal bovine serum (FBS; Wako) and 1% penicillin–streptomycin (Nacalai Tesque) at 37°C under 5% $CO_2$.

#### Primary neuronal culture

Primary hippocampal neurons were obtained from embryonic day 21 Sprague-Dawley rats. The embryos were decapitated, and the hippocampi were dissected, finely minced, and then treated with trypsin (Thermo Fisher Scientific) and DNase I (Sigma-Aldrich). The dissociated neuronal cells were plated onto glial cell monolayers cultured on a coverslip (Matsunami glass) coated with laminin (Thermo Fisher Scientific) and poly-L-lysine (Nacalai Tesque). Neuronal cultures were maintained in Neurobasal A medium supplemented with B-27 supplement (Thermo Fisher Scientific), 0.5 mM Glutamax (Thermo Fisher Scientific), 1 mM sodium pyruvate (Nacalai Tesque), and 1% penicillin/streptomycin (Nacalai Tesque) at 37°C under 5% $CO_2$. Cytosine $\beta$-D-arabinofuranoside (Sigma-Aldrich) was added to the culture medium, to a final concentration of 2.5–5 $\mu$M, at 2 days *in vitro*.

### Cloning and Plasmid Constructs

Recombinant DNAs for ELKS2/CAST, RIM1, RIMBP2, and Liprin-$\alpha$3 were PCR-amplified from a rat brain cDNA library (Thermo Fisher Scientific) and cloned into pHR lentiviral backbones or pcDNA3.1. In pcDNA3.1, ELKS2 was fused N-terminally to Lyn1-myristoylation and SNAP-tag or EGFP; RIM1$\alpha$ was C-terminally tagged with EGFP, SNAP, or Halo plus a KRas CAAX motif; covalent SNAP–ELKS2–RIM1(ΔN) was generated by overlap-extension PCR; RIMBP2 and Liprin-$\alpha$3 were tagged with EGFP at either terminus. Constructs were transformed into Stellar cells (Clontech), single colonies were selected and grown in LB medium with 100 $\mu$g/mL ampicillin at 37°C for 16h, and plasmid DNA was purified using the QIAprep Spin Miniprep Kit (Qiagen) according to the manufacturer's instructions. All constructs were validated by Sanger sequencing (GENEWIZ). The list of constructed plasmids is summarized in Table S1.

### Lentiviral transduction

Lentiviral particles for U2OS cells were generated by co-transfecting Lenti-X 293T cells grown to approximately 70% confluency, controlled with Countess 3 FL (Thermo), in 6-well plates with transfer plasmids, pCMV-dR8.91 and pMD2.G using FuGENE HD Transfection Reagent (Promega) following the manufacturer's instruction. After two days, 2 mL of supernatant containing viral particles was harvested and filtered with a 0.45 $\mu$m filter (Merck), and stored at –80°C until use. For primary hippocampal neuronal cultures, HEK 293T cells were co-transfected with transfer vector encoding EGFP–ELKS2 under the mouse CaMKII promoter, psPAX2 (Addgene), and pMD2.G (Addgene) using X-tremeGENE HP (Sigma-Aldrich). At 24h post-transfection, the medium was replaced with Neurobasal A medium, and viral supernatant was collected at 48h, and stored at –80°C.

### Construction of stable cell lines

For U2OS cells seeded at 10-20% confluency in 96-well glass-bottom plates (Cellvis), lentiviral supernatant was added directly to the culture medium. After 48h, the virus-containing medium was replaced with fresh growth medium, and cells were imaged no earlier than 72h post-infection. For primary hippocampal neuronal cultures, lentiviral particles were added at 7 days *in vitro* (DIV7), and neurons were maintained for at least an additional 7 days before immunocytochemistry and imaging.

**STED Imaging of In Situ Active Zones**

Immunohistochemistry of ELKS2 and RIM1 in brain sections was performed as previously described *(39)*. Briefly, Sprague-Dawley rats of either sex (postnatal day 44–46) were deeply anesthetized with isoflurane and quickly decapitated. The dissected brains were transferred into Tissue-Tek Cryomolds (Sakura Finetek), immersed in FSC 22 Frozen Section Media (Leica), and instantaneously frozen on an aluminum block in liquid nitrogen. Frozen brains were stored in liquid nitrogen until use. For sectioning, frozen brains were placed in a cryostat (CM1860, Leica) at −18°C for 30 min. Then, 10 $\mu$m-thick brain cryosections containing somatosensory cortex were sliced and retrieved on coverslips (Matsunami glass, 25 × 25, no. 1, 0.13 to 0.17 mm). The retrieved sections were immediately fixed by dehydration with a heat blower for 1 min. Sections were further fixed with ethanol at −25°C for 30 min and with acetone for 10 min on ice. After rehydration with PBS, sections were incubated with 0.3% BSA (Nacalai Tesque) containing PBS (blocking solution) for 30 min. Antibodies were diluted with the blocking solution and reacted with the preparations. Primary antibodies were applied for 3 hours at room temperature (RT) and washed twice with the blocking solution. Secondary antibodies were applied for an hour at RT, and then washed twice with the blocking solution and twice with PBS. The immunostained sections were lastly postfixed with 4% paraformaldehyde (PFA, FUJIFILM Wako Pure Chemical) for 15 min at RT and mounted in ProLong™ Glass Antifade Mountant (Thermo Fisher Scientific). Primary antibodies: rabbit polyclonal anti-ELKS2/CAST (2.5 $\mu$g/mL) *(39)*; mouse monoclonal anti-RIM1 (2.5 $\mu$g/mL; Synaptic Systems, Cat.No.140 111, IgG1, Clone 360H6E4, RRID:AB2864790). Secondary antibodies: donkey anti-rabbit IgG (H+L), highly cross-adsorbed (Jackson ImmunoResearch, Cat.No. 711-005-152) conjugated to STAR635P NHS Ester (Abberior); goat anti-mouse IgG1 (Jackson ImmunoResearch, Cat.No. 115-005-205) conjugated to Alexa Fluor 555 NHS Ester (Thermo Fisher Scientific).

Wide field brain imaging (Fig. 1d) was performed on a keyence BZ-700 all-in-one fluorescence microscope equipped with a metal halide lamp. The RFP and Cy5 excitation/emission filter set were used for Alexa Fluor 555 and STAR635P, respectively. The exposure time was set to 0.5 s and

1.0 s, respectively. Multiple images covering the entire brain were acquired with a 10× objective, and then stitched using BZ-X Analyzer software (Keyence, Japan).

STED imaging experiments were performed on a Leica TCS SP8 STED 3x microscope equipped with a pulsed white light laser, a continuous 592 nm STED laser, a pulsed 775 nm STED laser, HyD-SMD detectors, and a 100× oil immersion objective (NA = 1.40). Excitation laser wavelength was set to 561 nm for Alexa Fluor™ 633 nm for STAR635P. Laser power for excitation and depletion was set to 25% and 75%, respectively. Alignment of the STED depletion laser was performed every 30 min. Images were acquired with 1024×1024-pixel resolution at 400 Hz. The optical zoom factor was set to 11.36, resulting in a pixel size of 10.0 nm. The detector was configured to counting mode with a gating from 0.5 to 6.5 ns. Confocal images of the same field of view were acquired prior to STED imaging for comparison.

### Live-Cell STED Imaging of Reconstituted Active Zones

COS-7 cells were transfected with SNAP-tag fusion plasmids using X-tremeGENE HP (Sigma-Aldrich) in 24-well plates (Corning #353047) coated with 0.01 % collagen (FUJIFILM Wako) two days before imaging. At 24 h post-transfection, cells were replated onto coverslips coated with 0.01 % collagen and poly-L-lysine (Nacalai Tesque). SNAP-tagged proteins were labeled with SNAP-Cell 647-SiR (NEB #S9102S) one day later; HaloTag fusions were labeled with HaloTag TMR ligand (Promega) as needed.

Live-cell STED imaging was performed on a Leica TCS SP8 STED 3× microscope under the same settings as fixed-cell experiments. Excitation wavelengths were 488 nm (EGFP), 561 nm (TMR), and 633 nm (SiR); depletion power was 25 %. Laser power for snapshots was 50 %, and for time-lapse imaging 5 %. STED alignment was checked every 30 min. Images (512×512 pixels) were acquired at a pixel size of 10 nm. Confocal images of the same field were collected for comparison.

### Live-Cell Imaging of ELKS2 and RIM1 condensates

Fluorescence images for partition cofficient measurements were taken using a spinning-disk confocal microscope (Yokokgawa CSU-W1) with Nikon 100x oil immersion objective (CFI Plan Apo

$\lambda$D, NA 1.45) and an Andor iXon Ultra 888 EMCCD camera on a Nikon Eclipse Ti2-E body. Samples were maintained at 37°C and 5% CO2 with a stage top incubator (Tokai Hit). 488 and 561 lasers (Stradus) were used for imaging mGFP and mCherry/mRuby3, respectively. All image acquisition was performed using Nikon NIS-Elements AR software.

#### STED Imaging of Primary Neuronal Active Zones

For STED imaging of ELKS2 in control versus EGFP-ELKS2–overexpressing primary neurons (Fig. 3h), neurons were first infected at DIV7 with lentivirus encoding EGFP-ELKS2 under the CaMKII promoter or left uninfected (control) and maintained for 7 days. An optimized immunocytochemical protocol *(3)* was then applied to label AZ proteins. Briefly, neurons were fixed with 1% PFA in PBS for 15 min at RT, and then permeabilized with 0.1% Triton-X in PBS for 10 min at RT, followed by three washes in PBS. After blocking with 0.3% BSA containing PBS, Primary antibodies diluted in blocking solution were applied for 1 h at RT. After washing with blocking solution, the specimens were incubated with secondary antibodies for 30 min at RT, followed by three washes with PBS. Finally, after extensive washing, specimens were post-fixed with 4% PFA at RT for 15 min and mounted in ProLong™ Glass Antifade Mountant (Thermo Fisher Scientific). Primary antibodies were rabbit polyclonal anti-ELKS2/CAST (2.5 $\mu$g/mL, *(39)*), mouse monoclonal anti-RIM1 (2.5 $\mu$g/mL, Synaptic Systems, Cat.No.140 111, IgG1, Clone 360H6E4, RRID:AB2864790), and mouse monoclonal anti-GFP (1.0 $\mu$g/ml; FUJIFILM Wako Pure Chemical, Cat# 012-20461, clone: mFX73, IgG2a, RRID: AB_664697). Secondary antibodies were highly cross-adsorbed donkey anti-rabbit IgG (H+L) labeled with STAR635P NHS Ester, goat anti-mouse IgG1 labeled with Alexa Fluor™ 555 NHS, and goat anti-mouse IgG2a (Jackson ImmunoResearch, cat#115-005-206) labeled with Alexa Fluor™ 488 NHS Ester (Thermo Fisher Scientific), respectively. Synaptic sites were identified by immunolabeling of ELKS2 and RIM1, and ELKS2-overexpressing synapses were identified by GFP signals.

### Glutamate Imaging

Glutamate imaging experiments were conducted as described previously *(3)*. Hippocampal neurons were incubated with an optical glutamate sensor complex comprising streptavidin, biotinylated eEOS, and biotinylated BoNT/C-HC in HBS buffer (25 mM HEPES, 125 mM NaCl, 2.5 mM KCl, 2 mM $CaCl_2$, 1 mM $MgCl_2$, 25 mM D-glucose, 10 mM ascorbate, 3 mM myo-inositol, and 2 mM sodium pyruvate, pH 7.4). Neurons were imaged after incubation using an Olympus IX-71 inverted microscope with a ×100 oil immersion objective (NA = 1.40), equipped with an EMCCD camera (iXon, Andor) and U-MGFPHQ filter set (Olympus). Fluorescence responses ($\Delta F/F_0$) were quantified in ImageJ, and synaptic sites were identified by post hoc immunocytochemistry for Bassoon (mouse monoclonal, 0.5 $\mu$g/mL; Enzo Life Sciences, clone SAP7F407, IgG2a, RRID:AB 2038857) and RIMBP2 (mouse monoclonal, 5.0 $\mu$g/mL; RRID: AB 2864790) *(39)*, using Alexa 555- and Alexa 647-conjugated goat anti-mouse IgG2a and IgG2b secondary antibodies, respectively (Jackson ImmunoResearch).

### Nanocluster Size Analysis

To measure the size of ELKS2 nanoclusters (Fig. 1g,k), STED images of ELKS2 (pixel size of 10 nm) were processed by unsharp masking to detect the edges of clusters. In this process, STED images filtered by gaussian blur with a radius of 5 pixels (50 nm) were subtracted from STED images filtered by gaussian blur with a radius of 3 pixels (30 nm). The resultant subtracted images were then binarized, and components larger than 10 pixels (0.001 µm$^2$) were included in the analysis. The obtained masks were used to quantify the size of clusters. Data analysis was performed using Mathematica software version 13.0.1 (Wolfram), and histograms were plotted using MATLAB.

### Protein sequence feature

The coiled-coil domains of ELKS2 were predicted from its amino acid sequences using MARCOIL (https://bcf.sib.swiss/Delorenzi/Marcoil/index.html). The original Marcoil program was downloaded and modified to ensure compatibility with a modern C++ compiler, and the adapted version was used for analysis. The scores of coiled-coil for each residue were moving averaged for 3 residues, and regions with a continuous score of 0.5 or higher were considered as coiled-coil domains. The IDRs of RIM1 were predicted from its amino acid sequences using three prediction algorisms, IUPred3, PrDOS, and PONDR (https://iupred3.elte.hu/, https://prdos.hgc.jp/cgi-bin/top.cgi, https://www.pondr.com/). The scores of IDRs for each residue were averaged for the three predictors, and moving averaged for 20 residues. Amino acid sequences with score of 0.5 or higher were considered as IDRs.

### AlphaFold2 structure prediction

The structures of a rat ELKS2 protein (residues 1–973) dimer (Fig. 4a) and a mouse RIM1 protein (residues 1-1463) monomer (Fig. S3) were predicted using AlphaFold2. The predictions were run using the AlphaFold-Multimer with MMseqs2 on google colab*(19)*.

### 2D Radial Distribution Analysis

To quantitatively evaluate the periodic spacing of ELKS2 and RIM1 supramolecular nanostructures, we calculated the two-dimensional radial distribution function $g(r)$ (Fig. 1l). Analyses were performed in MATLAB (MathWorks) using the “2D Radial Distribution Function” script (`https://jp.mathworks.com/matlabcentral/fileexchange/132153-2d-radial-distribution-function`). The function is defined as

$$g(r) = \frac{1}{2\pi\, r\, \Delta r\, \rho\, N} \sum_{i=1}^{N} n_i(r, \Delta r), \tag{S1}$$

where $r$ is the distance from a reference particle, $\Delta r$ is the bin width, $\rho$ is the mean particle density, $N$ is the number of reference particles, and $n_i(r, \Delta r)$ is the number of other particles located within $r \pm \Delta r/2$ of reference particle $i$. We used a bin width of 1 px (10 nm) and computed $g(r)$ up to a maximum radius of 150 px in 512×512-px images to extract the characteristic nanostructure spacing.

#### Intensity Profile Analysis

Dot and ring nanostructures shown in Fig. 2a were quantified by drawing straight line ROIs across individual condensates in ImageJ (NIH) and extracting fluorescence intensity along those lines. The resulting intensity vs. distance profiles were exported and plotted in MATLAB, where peak-to-peak spacings were measured to determine the periodicity of ELKS2 dots and RIM1 rings.

#### Partition Coefficient Measurement

As shown in Fig. 2g, the partition coefficient $K$ for ELKS2 or RIM1 condensates was defined as

$$K = \frac{I_{\mathrm{cond}}}{I_{\mathrm{dil}}}, \tag{S2}$$

where $I_{\mathrm{cond}}$ and $I_{\mathrm{dil}}$ are the mean fluorescence intensities of the protein of interest in the condensate (dense phase) and the surrounding dilute phase, respectively. Condensate boundaries were identified by automated segmentation of the Gaussian-filtered EGFP or mRuby3 channel in MATLAB. To avoid interface artifacts, segmented condensates were eroded by three pixels ($\approx 0.48\,\mu$m), and intensity measurements were performed on the resulting regions. Gibbs free energy changes were calculated as

$$\Delta G = -k_{\mathrm{B}} T \ln K, \tag{S3}$$

where $k_{\mathrm{B}}$ is the Boltzmann constant and $T$ the absolute temperature.

#### Numerical Simulation of Synaptic Active Zones

Interpreting ELKS2–RIM1 supramolecular assemblies as a diblock copolymer (Fig. 3a), we modeled their microphase separation using the Ohta–Kawasaki free energy functional *(17)*:

$$\begin{aligned} F = \frac{k_{\mathrm{B}} T}{a^2} \int \Big[ & f_0(\phi) + \tfrac{\xi_0^2}{2} (\nabla\phi)^2 \\ & + \alpha\, G(\mathbf{r} - \mathbf{r}') \big(\phi(\mathbf{r}) - f_{\mathrm{ELKS2}}\big) \big(\phi(\mathbf{r}') - f_{\mathrm{ELKS2}}\big) \Big]\, d\mathbf{r}\, d\mathbf{r}', \end{aligned} \tag{S4}$$

$$\alpha = \frac{9}{2\,N^2\,a^2\,f_{\text{ELKS2}}^2\,(1 - f_{\text{ELKS2}})^2}, \tag{S5}$$

$$f_0(\phi) = \frac{a_2}{2}\,\phi^2 \;+\; \frac{a_4}{4}\,\phi^4, \tag{S6}$$

where $F$ is the Ginzburg-Landau free energy, $a$ denotes the unit cell length, $N$ is the total monomer number, $\xi_0$ is the correlation length, $G$ is the Green function, $f_{\text{ELKS2}}$ is the volume fraction of ELKS2, $a_2$ and $a_4$ are the negative and positive coefficients, respectively.

Phase separation dynamics can be described using Model B *(40)*:

$$\begin{aligned} \frac{1}{L}\,\frac{\partial\phi}{\partial t} &= \nabla^2\big(-\phi + \phi^3 - \nabla^2\phi\big) \\ &\quad - \frac{3}{16\,(4 - \chi N)^2\,f_{\text{ELKS2}}^2\,(1 - f_{\text{ELKS2}})^2}\big(\phi - f_{\text{ELKS2}}\big), \end{aligned} \tag{S7}$$

where $L$ is the transport coefficient.

The Flory parameter $\chi$ is defined by

$$\chi = \frac{z}{k_{\text{B}}T}\Big[\omega_{ER} \;-\; \tfrac{1}{2}(\omega_{EE} + \omega_{RR})\Big], \tag{S8}$$

where $\omega_{EE}$, $\omega_{RR}$, and $\omega_{ER}$ denote interaction energies of ELKS2–ELKS2, RIM1–RIM1, and ELKS2–RIM1 pairs, respectively, and $z$ is the coordination number. For $\chi > \chi_c \approx 7.3$, energetic penalties overcome mixing entropy, driving microphase separation into hexagonal, lamellar, and coexisting nanodomains as a function of $f_{\text{ELKS2}}$. Although this framework is derived in three dimensions, the final numerical simulations were implemented in two dimensions to directly compare with experimental membrane-anchored condensates.

### Pairwise MSD Analysis

To quantify collective spacing dynamics of ELKS2–RIM1 supramolecular particles (Fig. 4c), we computed the mean squared centroid distance—here referred to as "pairwise MSD"—for all particle pairs. Inter-particle trajectories $(x_j(t), y_j(t))$ and $(x_k(t), y_k(t))$ were extracted from microscopy images and processed in MATLAB (MathWorks) using custom scripts. For each pair $(j, k)$, the inter-particle distance

$$d_{jk}(t) = \sqrt{[x_j(t) - x_k(t)]^2 + [y_j(t) - y_k(t)]^2} \tag{S9}$$

was computed, and the pairwise MSD at time lag $\Delta t$ was defined as

$$\mathrm{MSD_{pair}}(\Delta t) = \left\langle [\, d_{jk}(t + \Delta t) - d_{jk}(t) \,]^2 \right\rangle_{t,(j,k)}, \tag{S10}$$

where $\langle\cdot\rangle_t$ denotes the time average over $t$ and $\langle\cdot\rangle_{(j,k)}$ the average over all particle pairs. Standard deviations at each $\Delta t$ were also calculated to assess variability in spacing fluctuations.

### Bond Orientational Order ($\psi_n$) Analysis

To quantify local spatial ordering of ELKS2–RIM1 supramolecular particles (Fig. S1), we used the Python library 'freud' (`https://freud.readthedocs.io/en/v1.2.2/examples/module_intros/Order-HexOrderParameter.html`). Under periodic boundary conditions, we performed a Voronoi tessellation with 'freud.locality.Voronoi' to identify each particle's neighbors. From these neighbor lists, we computed the bond orientational order parameters $\psi_n$ for symmetry orders $n = 4, 5, 6, 7$. For each particle $p$,

$$\psi_n^{(p)} = \frac{1}{N_p} \sum_{j=1}^{N_p} \exp\big(i\, n\, \theta_{pj}\big), \tag{S11}$$

where $N_p$ is the number of neighbors of particle $p$ and $\theta_{pj}$ is the angle of the vector from particle $p$ to neighbor $j$. We then used $|\psi_n^{(p)}|$ to quantify the strength of $n$-fold local order.

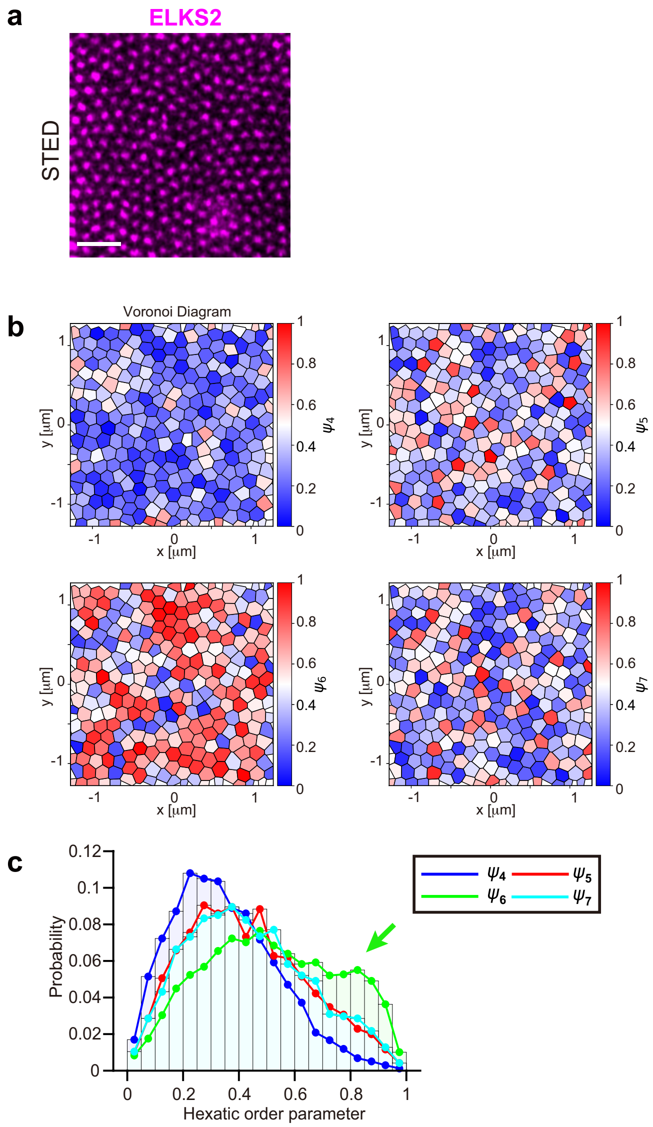


**Figure S1**: **Bond orientational order in ELKS2–RIM1 nanoclusters.** (a) STED image of a COS-7 cell expressing SNAP-ELKS2 and RIM1-EGFP. Scale bar: 500 nm. (b) Its Voronoi diagrams showing the hexatic order parameter $\psi_n$ for n=4, 5, 6 and 7, represented by their corresponding colors (see the method). (c) Probability distribution of various hexatic order parameters $\psi_n$ for n=4, 5, 6 and 7. The green arrow indicates the frequent emergence of six-fold symmetric structures. ($n_{\text{cell}}$, $n_{\text{dots}}$) = (7, 8019). $n_{\text{cell}}$: number of cells. $n_{\text{dots}}$: total number of dots.

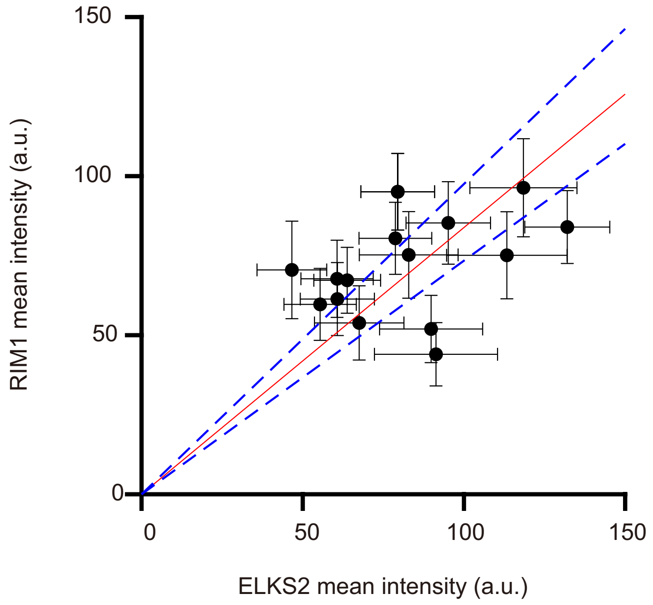


**Figure S2**: **Mean intensities of ELKS2 and RIM1 within condensates in non-neuronal COS-7 cells expressing SNAP-ELKS2(WT)-RIM1(ΔN)-EGFP, fused to a membrane translocation signal and localized to the plasma membrane, with each dot representing the mean intensity ± SD of individual condensates within a single cell.** A linear fit to the data points was performed to obtain the calibration line (red line), determining the fluorescence intensity ratio between SNAP-SiR and EGFP. The slope of the linear fit is 0.8385±0.01, and the dashed blue lines indicate the confidence interval of the fit. $n_{\text{cell}} = 16$

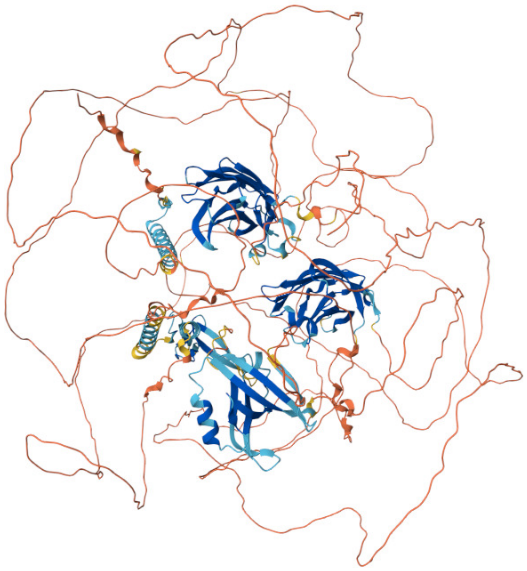

**Figure S3**: **AlphaFold2–predicted structure of RIM1, including the N-terminal zinc finger, PDZ domain, predicted central long intrinsically disordered region (IDR), and C-terminal C2B domain.**

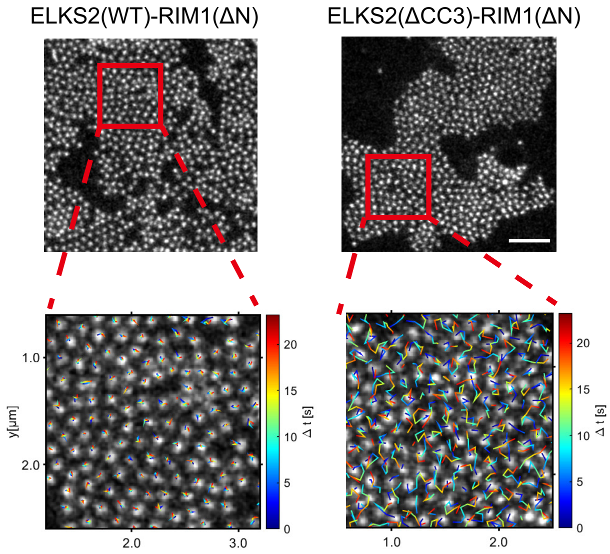


**Figure S4**: **Deletion of ELKS2's CC3 converts confined nanocluster motion into near-free diffusion. (a) STED images of COS-7 cells expressing SNAP–ELKS2–RIM1(ΔN)-EGFP with intact CC3 (WT, left) or CC3 deleted (ΔCC3, right).** Scale bars, 1 $\mu$m. Below each image, magnified regions (red boxes) show overlaid single-particle trajectories from 25 s time-lapse imaging, colored by elapsed time $\Delta t$ (0–25 s; color scale bar at right).

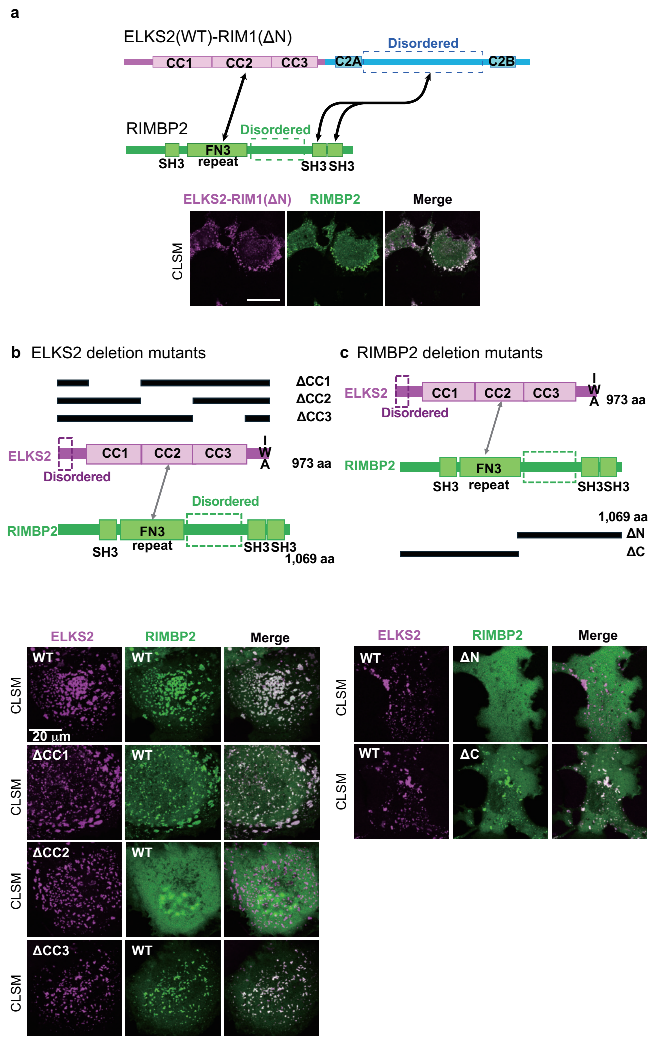


**Figure S5**: **Characterization of ELKS2–RIMBP2 interactions in reconstituted condensates.** (a) Top: Domain schematics of the covalent ELKS2–RIM1(ΔN) scaffold (magenta) and full-length RIMBP2 (green), showing CC1–CC3, disordered regions, FN3 repeat, and SH3 domains. Bottom: CLSM of COS-7 cells co-expressing SNAP–ELKS2–RIM1(ΔN)–Halo (magenta) with EGFP–RIMBP2 (green); merged signal (white) indicates co-localization in condensates. Scale bar: 20 $\mu$m. (b) Top: Domain maps of ELKS2 variants lacking individual coiled-coil domains CC1, CC2, or CC3 alongside RIMBP2 (green). Bottom: CLSM of each SNAP–ELKS2 truncation (magenta) co-expressed with EGFP–RIMBP2 (green); merged channels (white) show incorporation into condensates. Scale bar: 20 $\mu$m. (c) Top: Domain architectures of ELKS2 with RIMBP2 variants lacking N-terminal FN3 repeats (ΔN) or C-terminal SH3 domains (ΔC). Bottom: CLSM images of COS-7 cells co-expressing SNAP-ELKS2 (magenta) with each EGFP-RIMBP2 truncation (green); merged channels (white) show incorporation into condensates. Scale bar: 20 $\mu$m.

**Table S1**: **Key Resources Table**

| REAGENT or RESOURCE | SOURCE |
|---|---|
| **Figure 1** | |
| Plasmid: pcDNA Myr-SNAPf-ELKS2 | This paper |
| Plasmid: pcDNA RIM1-EGFP-CAAX | This paper |
| Plasmid: pcDNA RIM1-SNAPf-CAAX | This paper |
| Plasmid: pcDNA EGFP-RIMBP2 | This paper |
| Antibody: Anti-ELKS2 rabbit polyclonal antibody | Sakamoto et al., 2022 |
| Antibody: Anti-RIM1 mouse monoclonal antibody | SynapticSystems |
| **Figure 2** | |
| Plasmid: pcDNA Myr-SNAPf-ELKS2 | This paper |
| Plasmid: pcDNA RIM1-EGFP-CAAX | This paper |
| Plasmid: pcDNA RIM1-SNAPf-CAAX | This paper |
| Plasmid: pcDNA Myr-EGFP-ELKS2 | This paper |
| Plasmid: pHR-SFFV-RIM1-mRuby3 | This paper |
| Plasmid: pHR-SFFV-RIM1(ΔPDZ)-mRuby3 | This paper |
| Plasmid: pHR-SFFV-mGFP-ELKS2 | This paper |
| Plasmid: pHR-SFFV-mGFP-ELKS2(ΔIWA) | This paper |

| REAGENT or RESOURCE | SOURCE |
|---|---|
| **Figure 3** | |
| Plasmid: pcDNA SNAPf-ELKS2-RIM1(ΔN)-EGFP-CAAX | This paper |
| Plasmid: pcDNA SNAPf-ELKS2(CC1x2)-RIM1(ΔN)-EGFP-CAAX | This paper |
| Plasmid: pcDNA SNAPf-ELKS2(CC1x3)-RIM1(ΔN)-EGFP-CAAX | This paper |
| Plasmid: pcDNA SNAPf-ELKS2(CC1x4)-RIM1(ΔN)-EGFP-CAAX | This paper |
| Plasmid: pcDNA SNAPf-ELKS2-RIM1(ΔN, $\Delta IDR_1$)-EGFP-CAAX | This paper |
| Plasmid: pcDNA SNAPf-ELKS2-RIM1(ΔN, $\Delta IDR_2$)-EGFP-CAAX | This paper |
| Plasmid: pcDNA SNAPf-ELKS2-RIM1(ΔN, $\Delta IDR_3$)-EGFP-CAAX | This paper |
| Plasmid: pcDNA Myr-SNAPf-ELKS2 | This paper |
| Plasmid: pcDNA RIM1-EGFP-CAAX | This paper |
| Plasmid: pLenti-CaMKII EGFP-ELKS2 | This paper |
| Antibody: Anti-ELKS2 rabbit polyclonal antibody | Sakamoto et al., 2022 |
| Antibody: Anti-RIM1 mouse monoclonal antibody | SynapticSystems |
| Antibody: Anti-GFP mouse monoclonal antibody | FUJIFILM Wako Pure Chemcal |

| REAGENT or RESOURCE | SOURCE |
|---|---|
| **Figure 4** | |
| Plasmid: pcDNA SNAPf-ELKS2(ΔCC1)-RIM1(ΔN)-EGFP-CAAX | This paper |
| Plasmid: pcDNA SNAPf-ELKS2(ΔCC2)-RIM1(ΔN)-EGFP-CAAX | This paper |
| Plasmid: pcDNA SNAPf-ELKS2(ΔCC3)-RIM1(ΔN)-EGFP-CAAX | This paper |
| Plasmid: pcDNA SNAPf-ELKS2-RIM1(ΔN)-Halo-CAAX | This paper |
| Plasmid: pcDNA Liprin$\alpha$-3-EGFP | This paper |
| Plasmid: pcDNA EGFP-RIMBP2 | This paper |
| Plasmid: pcDNA EGFP-RIMBP2(ΔN) | This paper |
| Plasmid: pcDNA EGFP-RIMBP2(ΔC) | This paper |
| Antibody: Anti-Bassoon mouse monoclonal antibody | Enzo Life Sciences |
| Antibody: Anti-RIMBP2 rabbit polyclonal antibody | Sakamoto et al., 2022 |
| **Recombinant DNA** | |
| ELKS2/CAST | Rat brain cDNA |
| RIM1 | Rat brain cDNA |
| RIMBP2 | Rat brain cDNA |
| Liprin$\alpha$-3 | Rat brain cDNA |
